# Structure-seq2 probing of RNA structure upon amino acid starvation reveals both known and novel RNA switches in *Bacillus subtilis*

**DOI:** 10.1101/2020.04.20.051573

**Authors:** Laura E. Ritchey, David C. Tack, Helen Yakhnin, Elizabeth A. Jolley, Sarah M. Assmann, Philip C. Bevilacqua, Paul Babitzke

## Abstract

RNA structure influences numerous processes in all organisms. In bacteria, these processes include transcription termination and attenuation, small RNA and protein binding, translation initiation, and mRNA stability, and can be regulated via metabolite availability and other stresses. Here we use Structure-seq2 to probe the in vivo RNA structurome of *Bacillus subtilis* grown in the presence and absence of amino acids. Our results reveal that amino acid starvation results in lower overall dimethyl sulfate (DMS) reactivity of the transcriptome, indicating enhanced protection owing to protein binding or RNA structure. Starvation-induced changes in DMS reactivity correlated inversely with transcript abundance changes. This correlation was particularly pronounced in genes associated with the stringent response and CodY regulons, which are involved in adaptation to nutritional stress, suggesting that RNA structure contributes to transcript abundance change in regulons involved in amino acid metabolism. Structure-seq2 accurately reported on four known amino acid-responsive riboswitches: T-box, SAM, glycine, and lysine riboswitches. Additionally, we discovered a transcription attenuation mechanism that reduces *yfmG* expression when amino acids are added to the growth medium. We also found that translation of a leader peptide (YfmH) encoded just upstream of *yfmG* regulates *yfmG* expression. Our results are consistent with a model in which a slow rate of *yfmH* translation caused by limitation of the amino acids encoded in YfmH prevents transcription termination in the *yfmG* leader region by favoring formation of an overlapping antiterminator structure. This novel RNA switch offers a way to simultaneously monitor the levels of multiple amino acids.

## Introduction

Bacteria have the capacity to adapt to changing nutritional conditions via sweeping changes in gene expression. These regulatory events are tightly controlled such that various metabolic products are produced in sufficient quantities to sustain growth. Although regulation of transcription initiation plays a critical role in controlling gene expression, a variety of regulatory events that occur after transcription initiation play pivotal roles in modulating gene expression. Several of these post-transcriptional mechanisms rely on RNA structural features to regulate the level of transcription and translation. Some of these mechanisms involve binding of metabolites, proteins, and small RNAs (sRNAs) to the mRNA, leading to regulated transcription termination (attenuation and antitermination), translation initiation, and mRNA stability (Babitzke et al., 2009; Gollnick and Babitzke, 2002; Grundy and Henkin, 2004; Waters and Storz, 2009). RNA aptamers bind to a ligand (e.g. metabolite, RNA, or protein) to regulate gene expression through structural switching. These RNA switches change structure upon ligand binding and cause the refolding of neighboring nucleotides to alter transcription termination or ribosome binding. For instance, intrinsic transcription termination requires an RNA hairpin followed by a U-rich sequence. In several transcription attenuation mechanisms, formation of an antiterminator (AT) structure allows transcription readthrough by preventing formation of an overlapping terminator hairpin. Formation of the AT can in turn be prevented by an upstream anti-antiterminator (AAT) structure that overlaps the AT, and thus promotes formation of the terminator to cause transcription termination. An RNA aptamer involving these alternative structures allows control of transcription termination by altering the availability of the sequences involved and demonstrates the importance of RNA structure in bacterial gene regulation (Gollnick and Babitzke, 2002).

Recent advances in genome-wide RNA structure probing methods allow investigation of these essential features of a transcriptome in vivo. We developed Structure-seq2 (Ritchey et al., 2017), an improved version of the Structure-seq method (Ding et al., 2014) that probes all RNAs directly in cells such that the effects of the native physico-chemical environment are taken into account. The global set of RNA structures derived from this method is referred to as the “RNA structurome”. Other genome-wide RNA structure-probing methods have been used in bacteria, some of which focus on in vitro analysis of RNA structure with different conditions to identify RNA aptamers (Del Campo et al., 2015; Righetti et al., 2016; Tapsin et al., 2018). Recent in vivo studies on the model Gram-negative bacterium *Escherichia coli* compared RNA structure in vivo to that in vitro and to that with stalled ribosomes to relate RNA structure to translation efficiency, revealing that transcripts with more stable structures have a lower translation efficiency, although the causality is not clear (Burkhardt et al., 2017; Mustoe et al., 2018). Additionally, several highly abundant static RNA structural features were identified upstream of ribosomal protein coding genes that could play important biological roles (Mustoe et al., 2018). These studies made important contributions to global structural trends in *E. coli* and highlighted some important RNA structures. In the present study, we examined global RNA structural changes that occur in vivo due to amino acid starvation in the model Gram-positive organism *Bacillus subtilis*. These changes provide insight into how bacteria use RNA structure to respond to metabolite availability and nutritional stress.

The stringent response in *B. subtilis* is a global regulatory system that allows bacteria to respond to nutritional stress, including amino acid starvation, through the use of the alaramone (p)ppGpp (Eymann et al., 2001). The stringent response diverts cellular resources away from cellular growth and division and into the production of amino acids, as compared to the general stress response system that responds in a more nonspecific protective manner (Hecker and Volker, 2001; Voelker et al., 1995). The global proteome and transcriptome changes of the stringent response in *B. subtilis* have been studied (Eymann et al., 2002), but global changes to the RNA structurome have yet to be considered. Since RNA structure can change rapidly in response to the availability of metabolites, we examined RNA structurome changes that occur in response to amino acid starvation. We also compared the changes in the RNA structurome under this stress condition to known regulons in *B. subtilis*, as well as to known amino acid responsive RNA switches. After demonstrating the ability of Structure-seq2 to report on these features within *B. subtilis*, we used our data to predict unknown regulatory features. We identified several localized regions in which DMS reactivity of residues changed in response to amino acid limitation. One of these regions upstream of *yfmG* was explored in more detail and we found that *yfmG* is regulated by a transcription attenuation mechanism that responds to amino acid availability. This mechanism requires *yfmH*, a 23 amino acid leader peptide coding sequence just upstream of *yfmG*. Our results are consistent with a model in which a reduced rate of *yfmH* translation caused by amino acid limitation promotes transcription readthrough into the *yfmG* coding region.

## Results

### Structure-seq2 libraries

To obtain the in vivo genome-wide RNA structure landscape, *B. subtilis* was grown to mid-exponential phase in the presence of all amino acids (+AA), or in the absence of amino acids except for glutamine (-AA); glutamine is an important nitrogen source and was included to improve growth. DMS treatment in a single-hit kinetics range (Supplemental Fig. S1) was performed, which results in methylation of the Watson-Crick (WC) face of solvent-exposed A and C nucleobases. RNA was then extracted, depleted of rRNA, and subjected to Structure-seq2. In this method, reverse transcription with random hexamer primers results in cDNAs that terminate immediately before the site of DMS methylation. Additional library preparation steps allow for cDNA amplification and Illumina sequencing (Ritchey et al., 2019; Ritchey et al., 2017). After background subtraction using a library generated without DMS, solvent-exposed bases in reads that mapped to the transcriptome were identified. Comparison of DMS reactivity profiles +/- AA was used to assess RNA structural changes that occurred in response to amino acid starvation. Each condition (+/- AA and +/- DMS) was performed in biological triplicate. We generated a total of 287M Structure-Seq2 reads that passed initial filtering (Supplemental Table W.1). Aligning these high-quality reads against a custom transcriptome yielded a 77% average rate of mapping (Supplemental Table W.2). These mappings were then filtered to remove alignments with a mismatch on the first base pair or more than 3 total mismatches via StructureFold2 (Supplemental Table W.3) (Tack et al., 2018). The efficacy of our DMS treatment was confirmed by high A and C specificity (Supplemental Table W.4; Supplemental Fig. S2). When +DMS replicates of each growth condition were pooled, coverage was sufficient (greater than 1 reverse transcriptase [RT] stop per A or C in the transcript) (Ding et al., 2014;

Ritchey et al., 2019; Su et al., 2018; Tack et al., 2020; Waldron et al., 2019) to resolve 2602 (+AA) and 2600 (−AA) transcripts, with an overlap of 2333 transcripts. The RT stops of +DMS replicates of both +AA and −AA treatments were well-correlated for all resolvable transcripts within that condition (Supplemental Fig. S3).

### Transcriptome-wide trends

The calculated DMS reactivity patterns for each transcript and its three regions (5’UTR, coding sequence [CDS], and 3’UTR) were analyzed to determine global effects of amino acid starvation. The overall trend was decreased DMS reactivity upon amino acid starvation for the whole transcript as well as the individual regions, although certain transcripts increased (Fig. 1A-D; Supplemental Table T.2, D.1-D.4). These observations imply a more structured and/or protein-protected structurome under amino acid starvation conditions.

**Figure 1.**
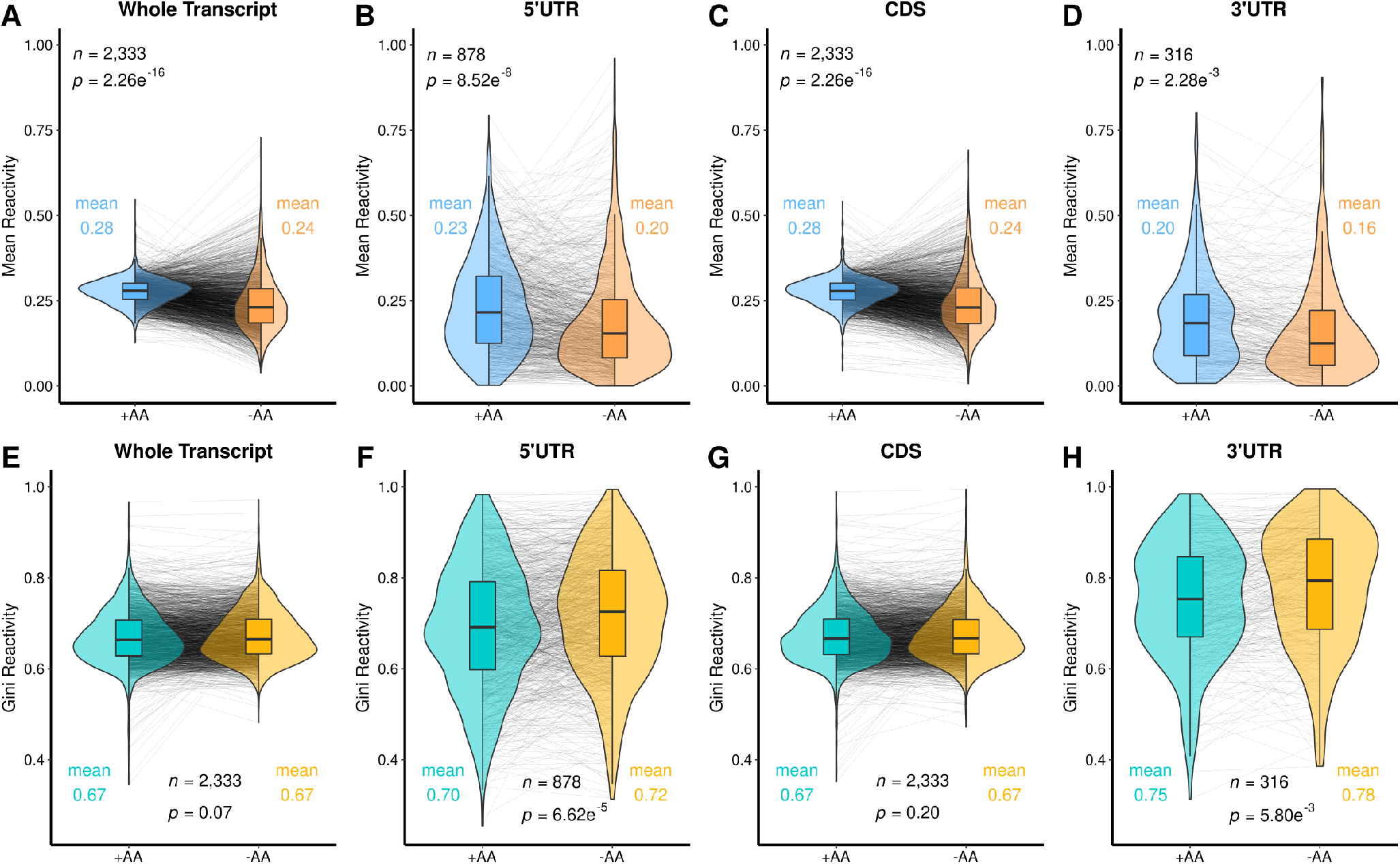
Amino acid starvation leads to lower DMS reactivity in all parts of transcripts, and increased Gini in the untranslated regions. (*A*) The distribution of mean transcript reactivities of the +AA transcriptome (teal) is compared to that of the −AA transcriptome (orange) on violin plots to show the overall distribution with a box plot at the center. Changes of individual transcripts are shown as gray lines. Most transcripts were less reactive when grown without amino acids. The overall means were compared via a t-test with the p-value reported in each panel. (*B-D*) 5’UTR, CDS, and 3’UTR respective regions following the same formatting as (*A*). (*E-H*) Mirrors of (*A-D*), but for the Gini coefficient of DMS reactivity rather than for the mean, between +/-AA transcriptomes. A table of these t-tests can be found in the Supplemental Table T.2

We also ascertained transcriptome-wide changes in the Gini index (Fig. 1E-H), as applied to DMS reactivity. Gini is a measure of the unevenness of the data, where a low Gini score has been found to be characteristic of unstructured regions of a transcript, as all nucleotides would have similar DMS reactivity, and a high Gini score reflects more structured regions (Rouskin et al., 2014). We found that the Gini score increased significantly in the 5’ and 3’UTRs in response to amino acid starvation (Fig. 1F and H; Supplemental Table T.2, D.2, D.4). Decreasing DMS reactivity coupled with increasing Gini index in the 5’ and 3’UTRs supports greater RNA structure and is generally consistent with reported lowered translation efficiency (Burkhardt et al., 2017), as expected from amino acid starvation.

### Change in DMS reactivity is inversely correlated with change in transcript abundance

In addition to structural information, Structure-seq2 provides transcript abundance, similar to RNA-seq (Su et al., 2018; Tack et al., 2018). Previously, we found that changes in transcript reactivity are inversely correlated with changes in transcript abundance following 10 min of heat shock in rice seedlings (Su et al., 2018) and after persistent salinity stress in Arabidopsis (Su et al., 2018; Tack et al., 2020). Those transcripts with increased DMS reactivity following heat shock or salt stress are thus less innately structured or protein protected and may have increased exposure to degradation pathways. On the other hand, those transcripts with decreased DMS reactivity following heat shock or salt stress may bind proteins or have increased RNA folding (salt stress only) to gain protection from such mechanisms (Su et al., 2018; Tack et al., 2018). We investigated changes in transcript reactivity with changes in transcript abundance (Supplemental Table D.5, D.6) for *B. subtilis* upon amino acid starvation, in part to see whether this trend is preserved in another domain of life.

We observed that globally, the change in mean transcript reactivity was inversely correlated with the change in abundance of that transcript (Fig. 2), just as was found in rice and Arabidopsis. Transcripts that exhibited increased DMS reactivity under amino acid starvation generally exhibited decreased abundance, whereas transcripts that exhibited decreased DMS reactivity in the absence of amino acids exhibited increased abundance (Fig. 2A; Supplemental Table T.3, D.1-D.6). This inverse relationship was observed when comparing whole transcripts as well as the CDS, while the UTR regions did not show this same inverse correlation (Fig. 2B-D).

**Figure 2.**
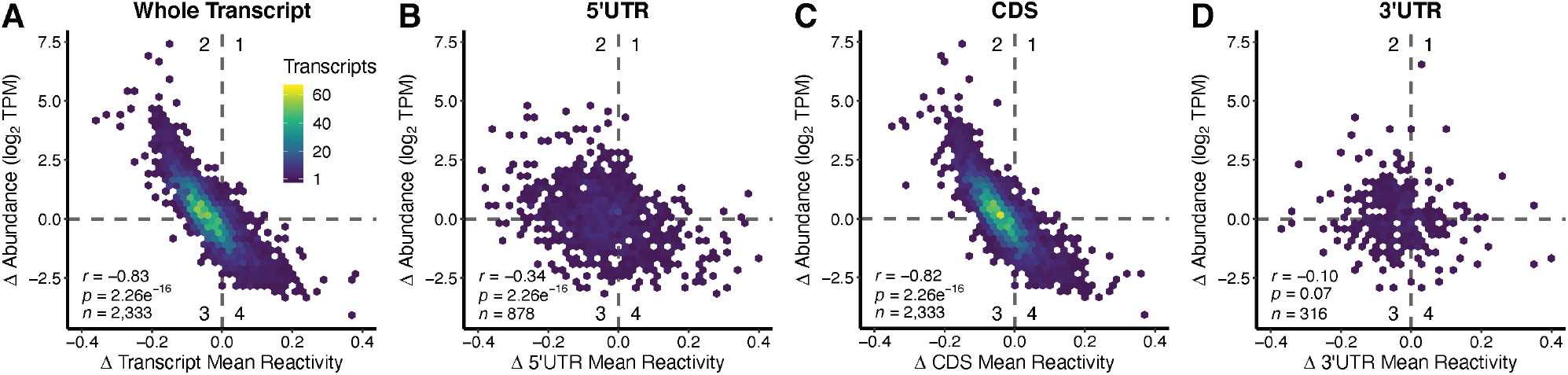
Change in transcript DMS reactivity and change in abundance are inversely correlated overall and in the CDS, but not in the 5’UTR or 3’UTR. (*A*) Change (Δ) in transcript mean DMS reactivity (mean DMS reactivity_-AA_ – mean DMS reactivity+AA) were plotted against change (Δ) in transcript abundance (log_2_(TPM_-AA_) - log_2_(TPM_+AA_)). These transcripts were then broken down into their component regions and the analysis repeated; (*B*) 5’UTR, (*C*) CDS, and (*D*) 3’UTR. A table summarizing all relevant tests and statistics for this figure can be found in Supplemental Table T.3.

Previous studies in the model plant species *Arabidopsis thaliana* showed that more reactive transcripts were more abundant within a given condition (i.e. comparing absolute amounts rather than changes in amount) (Li et al., 2012), so we sought to confirm if this was also a trend in *B. subtilis*. Indeed, transcripts with higher average DMS reactivity within a condition were more abundant (Fig. 3A). This is distinct from the inverse correlation between the *change* in DMS reactivity and the *change* in abundance. The reactivity and coverage for each individual structurome (+AA, −AA) was recalculated (Supplemental Fig. S4; Supplemental Table D.7, D.8), yielding more transcripts than the contrasted structuromes, as mutual coverage of transcripts between conditions was not considered, and less dispersed numbers, as the reactivity values were normalized by their respective internal 2-8 % normalization scales rather than cross normalized by the same 2-8% scale to ensure an accurate comparison between conditions. Similar trends show that in each separate structurome, the same relationship holds, where more reactive transcripts are typically more abundant (Supplemental Fig. S4). The trend is slightly stronger in −AA conditions (r = 0.43 vs. r = 0.67, Fisher Z-test p = 0.0000), again pointing to more overall modulation of structure in stress conditions. Additional details can be found in Supplemental Table T.4. We also plotted each of the transcripts as a straight-line arrow to show the shift that occurred when changing from growth in the presence of amino acids to growth in their absence (Fig. 3B). The majority of changes reciprocated each other: an increase of DMS reactivity corresponded with a decrease in abundance (red, sloping downwards), while a decrease in DMS reactivity was associated with an increase in abundance (blue, sloping upwards). This was highlighted by plotting just the 5% largest increases and decreases in abundance (Fig. 3C).

**Figure 3.**
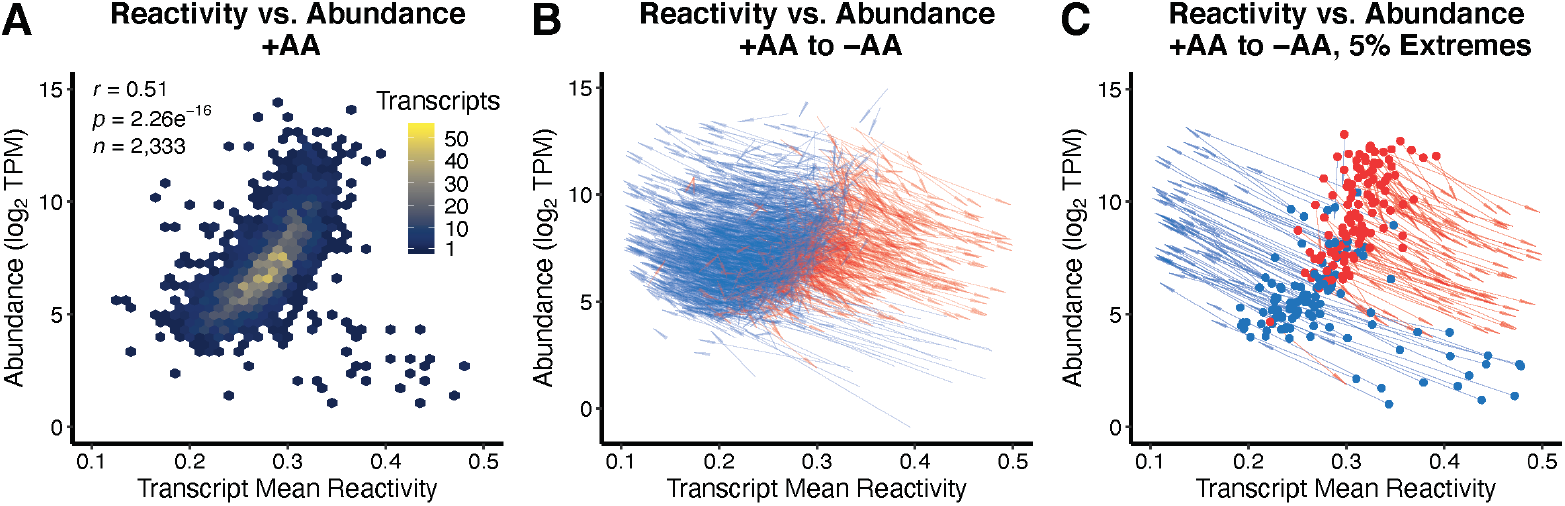
Average DMS reactivity positively correlates with transcript abundance. (*A*) The mean transcript reactivity is positively correlated to transcript abundance in +AA conditions, such that more reactive, accessible transcripts tend to be more abundant (r = 0.51, p < 2.26e^-16^). (*B*) All transcripts resolved in both +/-AA conditions are plotted as an arrow originating at the reactivity and abundance (log_2_ TPM) of each respective transcript’s +AA values, and ending at that transcripts −AA reactivity and abundance values. (*C*) A subset of (*B*), limiting the data to the top 5% largest increases and decreases of transcript abundance for better visualization. Arrow lines show where DMS reactivity decreased (blue) or increased (red) from +AA to −AA, while arrow origins show where abundance decreased (red) or increased (blue) from +AA to −AA.

### The stringent response and CodY regulons show coordinated changes in abundance and DMS reactivity in response to amino acid availability

Bacterial survival depends upon the capacity to rapidly alter metabolism, physiology, and behavior in response to changing environmental conditions. These responses involve changes in the expression of numerous genes. To coordinate these responses, bacteria utilize global regulatory networks through the action of a limited repertoire of regulatory factors (Cavanagh and Wassarman, 2014; Eymann et al., 2002; Fujita, 2009; Geiger and Wolz, 2014; Kovacs, 2016; Romeo et al., 2013; Sonenshein, 2005). As RNA abundance is a common way in which cells regulate metabolic pathways in response to certain signals, we sought to determine if a change in RNA accessibility also related to metabolic control. To determine how these RNA structural trends may affect physiology, we considered regulons, which are defined as sets of genes that are controlled by the same signal or regulatory molecule. We sought to uncover whether RNA structural features may contribute to coordination of these regulons. After assessing 174 known regulons in *B. subtilis* for a relationship between the changes in DMS reactivity and abundance as obtained through Structure-seq2 (Supplemental Item 1), we found that transcripts encoded by genes in both the stringent response and the CodY regulons experienced pronounced coordinated changes in abundance and DMS reactivity in response to amino acid availability (Fig. 4).

The stringent response regulon is the set of genes that are positively or negatively regulated in response to the alarmone (p)ppGpp following nutritional stress, including amino acid starvation (Eymann et al., 2002). Transcripts corresponding to genes that are known to be positively regulated by the stringent response showed a decrease in DMS reactivity and we confirmed their previously reported increased abundance (Eymann et al., 2002) when grown in the absence of amino acids, while genes known to be negatively regulated showed an increase in DMS reactivity, and we confirmed their previously reported decreased abundance (Eymann et al., 2002) (Fig. 4A-C; Supplemental Table L.1). These results indicated that the inverse relationship has functional consequences for the expression of genes that are critical for the ability of *B. subtilis* to survive amino acid starvation, controlling abundance through nucleotide accessibility. Nearly all of the transcripts positively regulated by the stringent response decreased in DMS reactivity, while nearly all of the negatively regulated transcripts increased in DMS reactivity.

**Figure 4.**
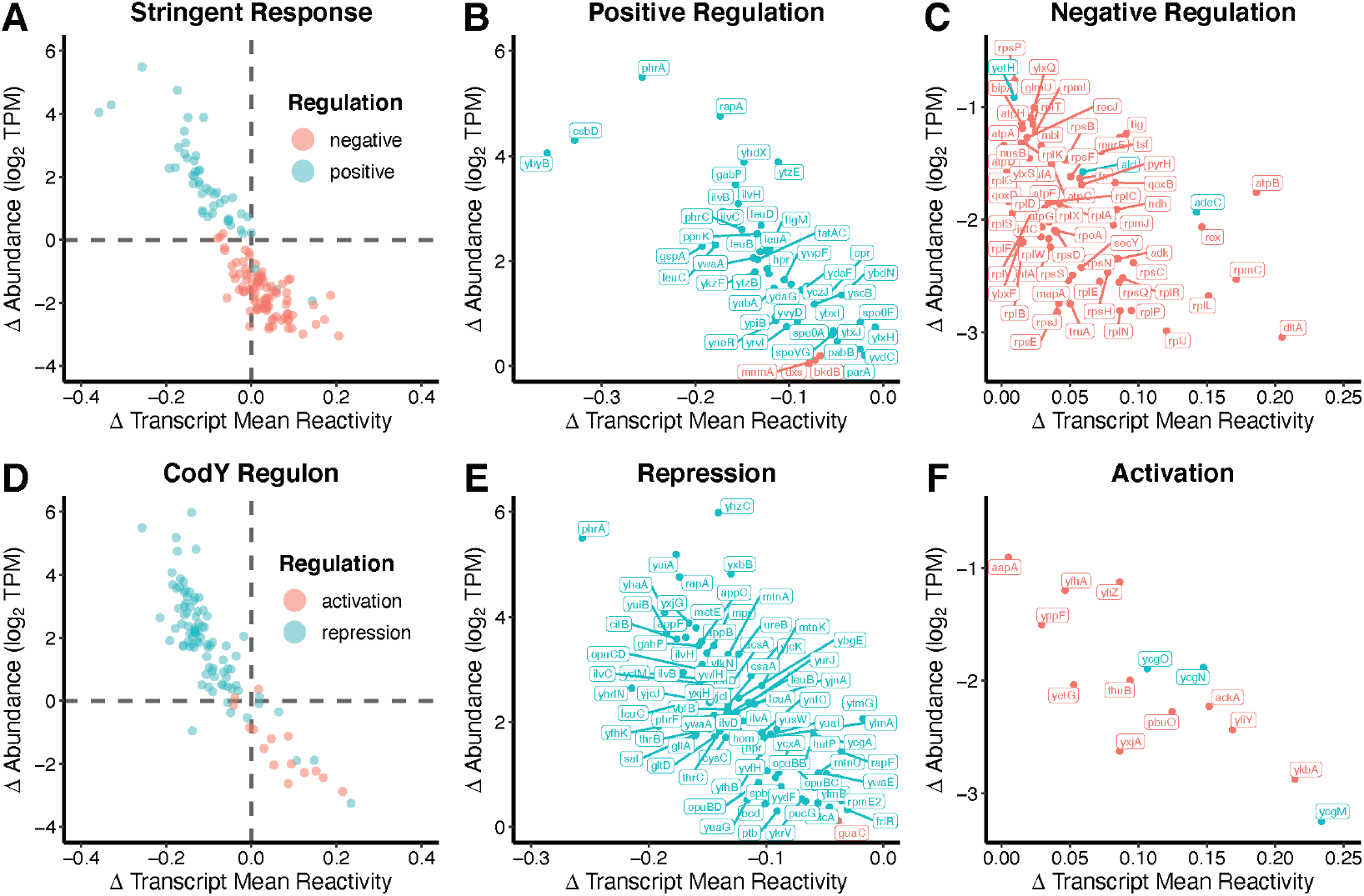
Transcript DMS reactivity and abundance changes in the stringent response and CodY regulons are controlled by RNA accessibility. (*A*) Transcripts previously reported to be positively (teal) and negatively (red) regulated by the stringent response were subset from the data and plotted, comparing the change (Δ) in reactivity against the change (Δ) in abundance as per Figure 2. Abundance changes from our work clearly parallel reported abundance changes; the majority of positively regulated transcripts increased in abundance in −AA media (quadrant II). Likewise, the majority of negatively regulated transcripts decreased in abundance (quadrant IV). (*B*) Transcript names of positively regulated transcripts from quadrant II and (*C*) negatively regulated transcripts from quadrant IV. (*D-F*) Same analysis but for the CodY regulon, where repressed transcripts are also found in quadrant II and activated transcripts in quadrant IV. A full table summarizing these data is included (Supplemental Table L.1, L.2).

The opposite trend was observed for the CodY regulon, which includes genes involved in adaptation to nutritional stress (Fig. 4D-F; Supplemental Table L.2). Branched chain amino acids and GTP serve as ligands for CodY, a DNA binding regulatory protein. The CodY regulon is linked to the stringent response because (p)ppGpp synthesis reduces GTP levels (Geiger and Wolz, 2014). Genes known to be activated by CodY showed an increase in DMS reactivity and we confirmed their previously reported decrease in abundance (Brinsmade et al., 2014), while repressed genes showed a decrease in DMS reactivity and we confirmed their previously reported increase in abundance (Brinsmade et al., 2014) when grown in the absence of amino acids. The other 172 regulons analyzed did not show this strong correlation between known levels of regulation and DMS reactivity changes, which underscores the specificity of the response to the stringent response and CodY regulons.

### Structure-seq2 identifies known riboswitches during changes to amino acid availability

The global RNA structural changes that we observed indicate that several mechanisms occur within the cell in response to amino acid availability. RNA switches are common mechanisms in which RNA structural features switch conformation following binding of another molecule. This switch then affects downstream gene expression. There have been numerous biochemical and molecular genetic studies characterizing various classes of RNA switches (McCown et al., 2017; Sherwood and Henkin, 2016). Each of these experimental studies has focused predominantly on one class of RNA switch at a time.

Currently, there are three known classes of direct amino acid-binding RNA switches: the lysine (Grundy et al., 2003; Sudarsan et al., 2003), glycine (Mandal et al., 2004), and glutamine (Ames and Breaker, 2011) riboswitches. Additionally, amino acids are sensed indirectly through the SAM riboswitches that sense methionine levels (McDaniel et al., 2003), the T-box elements that sense the amount of uncharged tRNAs (Green et al., 2010), and protein-responsive elements such as the tryptophan-responsive TRAP binding elements (Babitzke and Gollnick, 2001). These well-characterized RNA switches provide a starting point to benchmark Structure-seq2 and thereby hint that there may be additional amino acid-responsive RNA switches, especially given that there are 20 common amino acids and a large number of intermediates in the cell.

We initially focused on 31 RNA switches that are known to respond to amino acids in *B. subtilis* (Supplemental Table L.3). The 30 RNA switches able to be resolved in StructureFold included the lysine and glycine riboswitches that directly bind amino acids (Grundy et al., 2003; Mandal et al., 2004; Sudarsan et al., 2003), T-box elements that monitor the charged state of tRNAs (Green et al., 2010), S-box riboswitches that bind S-adenosylmethionine (McDaniel et al., 2003), and the protein-mediated RNA switches for tryptophan (Gollnick et al., 2005). The overall change in mean DMS reactivity of the 5’UTRs of the 30 targeted transcripts is not significant (Fig. 5A, p= 0.71). This is expected since ligand binding to riboswitches generally involves a trading of base pairs rather than any large net loss or gain. This issue is examined in detail below. While these 5’UTRs do change mean DMS reactivity between conditions, this overall change may not mirror or be indicative of changes in the switches and functional elements. The average DMS reactivity decreased in the CDS for these 30 targeted transcripts upon amino acid starvation, as seen with all transcripts (Fig. 1C, Fig. 5B, p = 3.25e-3).

**Figure 5.**
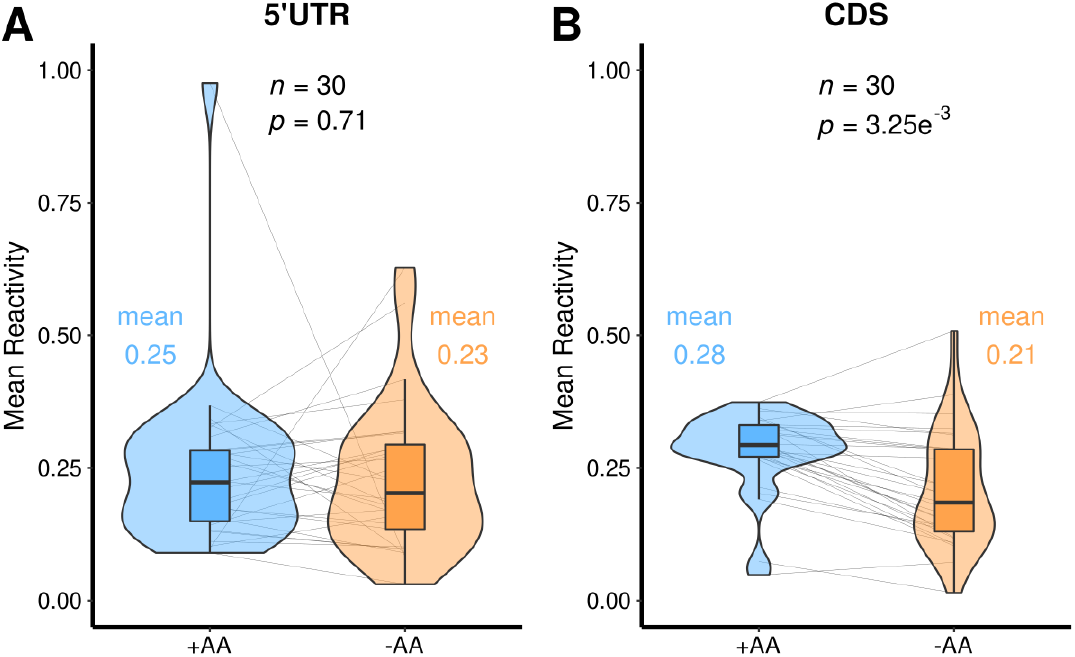
Violin plots of the 30 genes known to respond structurally to amino acids. (*A*) 5’UTR region. (*B*) CDS region. These data can be found in Supplemental Table D.9, D.10.

We plotted the change in DMS reactivity after amino acid starvation onto known secondary structures and crystal structures of some of the characterized RNA switches, including the T-box, S-box, glycine, and lysine riboswitches (Fig. 6; Supplemental Fig. S5; Supplemental Table L.4). Structure-seq2 faithfully reported on nucleotides known to be protected by the ligand, and the data aligned well with previously reported in vitro probing data in which changes in RNA structure were observed depending on the absence or presence of bound ligand.

**Figure 6.**
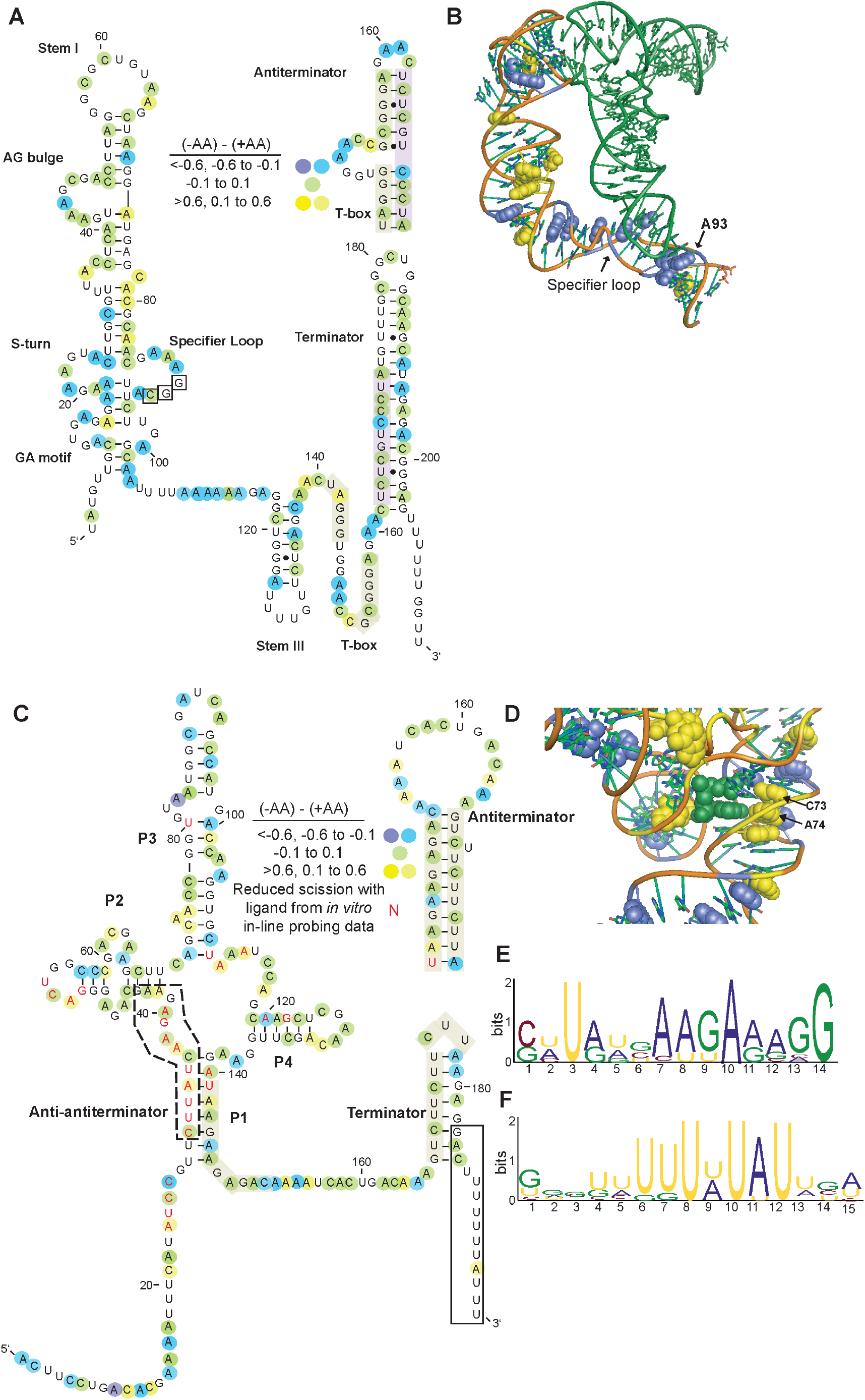
Structure-seq2 data align well with known riboswitches and DMS reactivity changes reveal functionally significant regions. (*A*) Change in DMS reactivity displayed on the secondary structure of the *B. subtilis glyQS* T-box element. Blue shows nucleotides that had decreased DMS reactivity without amino acids, green shows nucleotides that did not change in DMS reactivity, and yellow shows nucleotides that had increased DMS reactivity without amino acids. Darker yellow and blue indicate more extensive changes of DMS reactivity. Secondary structure model adapted from (Caserta et al., 2015). (*B*) Change in DMS reactivity displayed on the crystal structure of the *G. kaustophilus glyQS* T-box (Grigg and Ke, 2013). Blue and yellow are nucleotides that had decreased or increased DMS reactivity without amino acids, respectively, and the tRNA ligand is displayed in forest green. PDBID: 4MGN (*C*) Change in DMS reactivity displayed on the secondary structure of the *B. subtilis yitJ* SAM riboswitch. Color scheme is the same as in (*A*), and nucleotides reported to have decreased DMS reactivity with bound SAM in a previous in vitro study are in red font (Winkler et al., 2003). (*D*) Change in DMS reactivity displayed on the crystal structure of the *yitJ* SAM riboswitch (Lu et al., 2010). Blue and yellow are nucleotides that had decreased or increased DMS reactivity without amino acids, respectively, and the SAM ligand is displayed in forest green. PDBID: 4KQY. (*E-F*) Highly overrepresented sequences that increased in DMS reactivity without amino acids from sliding windows of the 5’UTRs of the 30 resolved transcripts. (*E*) This region corresponds to the anti-antiterminator which would open up upon removal of SAM (dashed box in *C*). (*F*) This region corresponds to the 3’-end of the terminator, which would become exposed upon removal of SAM (black box in *C*).

The T-box element is a structurally responsive element that binds uncharged tRNAs and is often found in the 5’UTR of genes encoding tRNA synthetases that charge the tRNA with the cognate amino acid. When uncharged tRNA binds to the nascent transcript it stabilizes the antiterminator conformation in the 5’UTR such that transcription continues into the downstream coding sequence, and the resultant increase in tRNA synthetase promotes the charging of tRNA molecules (Grundy and Henkin, 2004; Sherwood and Henkin, 2016). In our system, the absence of amino acids would drive tRNA to bind to the T-box riboswitch, presumably leading to protection from DMS reactivity. While there are multiple T-box riboswitches found within *B. subtilis*, the published secondary structure of the *glyQS* T-box was used to demonstrate structural changes associated with T-box functionality (Caserta et al., 2015). Although most of the nucleotides participating in the structure of the *glyQS* T-box did not show altered DMS reactivity values (Fig. 6A) (Caserta et al., 2015), there are key residues that were altered in the absence of amino acids. For instance, when plotting the change in Structure-seq2 DMS reactivity on the part of the *Geobacillus kaustophilus glyQS* T-box that has a solved crystal structure (Grigg and Ke, 2013), one nucleotide in the specifier loop, A93, is less reactive in the absence of amino acids, likely because it is protected by bound tRNA (Fig. 6B). When bound with tRNA, this nucleotide stacks on the wobble position of the anticodon and forms a hydrogen bond with a separate residue (Zhang and Ferre-D’Amare, 2013). Additionally, A89 is known to form a sheared AA base pair that is near the tRNA that could account for the lower reactivity when tRNA binds. This binding promotes the formation of a pocket to accommodate tRNA modifications and could also explain the lower reactivity of nearby A87 when tRNA binds in the absence of amino acids (Zhang and Ferre-D’Amare, 2013).

The S-box, or SAM riboswitch, binds S-adenosylmethionine, which contains methionine, and promotes termination (Grundy and Henkin, 1998). There were a substantial number of nucleotides within the SAM riboswitch of the *yitJ* mRNA in *B. subtilis* that changed DMS reactivity in the absence of amino acids (Fig. 6C) (McDaniel et al., 2003). When aligning published in vitro probing data with our in vivo Structure-seq2 data, nucleotides that had data in both datasets were in good agreement (Winkler et al., 2003). For instance, A24, A34, A37, A52, A111, and A113, which are ligand-protected in vitro, had increased DMS reactivity in the absence of amino acids in vivo (Fig. 6C, note overlap of red lettering with yellow shading). This increased DMS reactivity suggests that the RNA is bound by SAM in the presence of amino acids and unbound in their absence. According to the crystal structure of the *B. subtilis yitJ* SAM riboswitch (Lu et al., 2010), two key residues that participate in recognizing the SAM moiety, C73 and A74, were exposed in the absence of amino acids in our in vivo Structure-seq2 data (Fig. 6D). Other SAM-protected nucleotides in the in vitro dataset showed no significant change in DMS reactivity in vivo, with the exception of A120, which showed *decreased* DMS reactivity without amino acids in vivo; these discrepancies could reflect the influence of full flanking sequence present in vivo, cellular factors, or differences in probing reactivity (OH^-^ for in-line probing vs DMS in Structure-seq2). While the T-box and S-box riboswitches monitor the availability of amino acids indirectly, the glycine and lysine riboswitches bind amino acids directly. Similar analyses were performed on these riboswitches, which supported our Structure-seq2 data by faithfully monitoring loss of amino acid binding upon starvation (Supplemental Fig. S5).

To identify regions of significant structure change more broadly, we used StructureFold2 (Tack et al., 2018) to perform a sliding window analysis throughout the 5’UTRs of the 30 transcripts known to contain amino acid responsive RNA switches, looking at regions with increased DMS reactivity in the absence of amino acids (Supplemental Table L.5). The top 25% of windows with increased DMS reactivity in the absence of amino acids revealed overrepresented sequences corresponding closely to conserved regions within the S-box riboswitches (Fig. 6E, F). One region of the RNA, which is in a dashed boxed in Figure 6C with MEME in Figure 6E, is known to become partially sequestered in an anti-antiterminator structure when SAM binds and would be released in the absence of SAM binding. In addition, a second region of the RNA, which is in a solid boxed region in Figure 6C with MEME in Figure 6F, makes up the extreme 3’ end of the terminator hairpin that would become released in the absence of SAM binding. These sliding windows thus revealed important functional regions within RNA switches.

### Discovery of a transcription attenuation mechanism via Structure-seq2

In addition to further characterization of known amino-acid responsive RNA switches, we sought to use our Structure-seq2 data to discover new RNA switches. Computational analyses to discover classes of RNA switches have been based primarily on conservation of sequence, structure, and/or gene function of neighboring genes (Weinberg et al., 2017). This analysis pipeline has been modified to uncover increasingly rare RNA switches (Weinberg et al., 2017), but there are limitations to these methods. For instance, RNA switches belonging to just one type of bacteria or without a conserved sequence or known regulatory mechanism may be missed using this methodology. Thus, there is the potential that additional novel RNA switches exist, especially considering the vast number of molecules within a cell (Stav et al., 2019). Since known RNA switches have been found predominately within the 5’UTR of transcripts, we chose to first perform a search for new candidates within the 5’UTRs. We analyzed the 5’UTRs plus the first fifty nucleotides of the coding sequence from the Structure-seq2 reactivity files. We used the sliding window module from StructureFold2 (Tack et al., 2018) to analyze the top 5% of DMS reactivity changes within 20 nt windows, sliding by 10 nt intervals, in an effort to identify those that have the greatest change in DMS reactivity between our two growth conditions (i.e. ± amino acids). Many of the windows generated corresponded to regions of genes with unknown function (Supplemental Table L.6). One such gene, *yfmG*, contains a region with homology to the formylglycine-generating enzyme domain (Caspi et al., 2014). Members of this family convert cysteine to formylglycine post-translationally (Fig. 7A) (Carlson et al., 2008). Within this 5’UTR a NusA-dependent intrinsic transcription terminator was identified previously by Term-seq (Fig. 7 B, C) (Mondal et al., 2016). There were large changes in DMS reactivity upon amino acid starvation (Fig. 7D), with the largest change upstream of this terminator. The structures predicted using in vivo constraints in RNAStructure (Reuter and Mathews, 2010) were predicted to have more base-pairing interactions in the presence of amino acids (Fig. 7 B, C).

**Figure 7.**
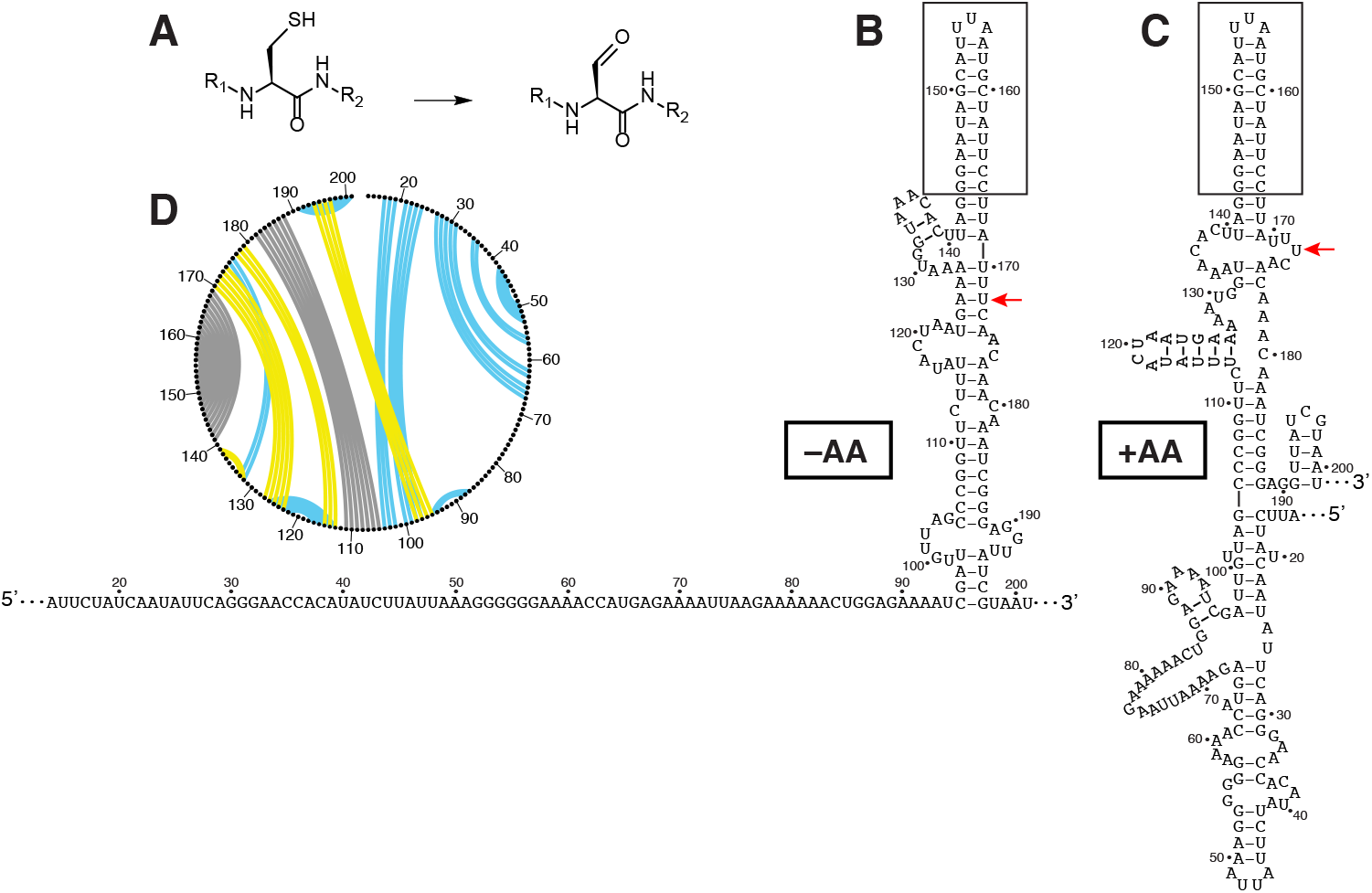
Amino acid responsiveness of the *yfmG* 5’UTR element. (*A*) Formylglycine generating enzymes catalyze the conversion of cysteine into formylglycine. (*B, C*), Secondary structures constrained with in vivo Structure-seq2 data in the absence (*B*), and presence (*C*) of amino acids. Notably, a known site of termination is shown as a red arrow with the terminator hairpin boxed (Mondal et al., 2016). (*D*) A CircleCompare plot showing changes in the RNA secondary structure from Structure-seq2 data upon amino acid starvation. Yellow lines are base pairs predicted only in the absence of amino acids, blue lines are base pairs predicted only in the presence of amino acids, and gray lines are shared base pairs.

The *yfmG* transcription start site was determined prior to conducting regulatory studies. Total cellular RNA was extracted from mid-exponential phase cultures grown under the same conditions used for Structure-seq2. Primer extension analysis led to the identification of a single *yfmG* transcription start site 201 nt upstream of the *yfmG* translation initiation codon (Fig. 8A, B), which allowed us to identify a σ^A^-dependent promoter with an extended −10 element (Fig. 8A). The long 201-nt leader suggested that *yfmG* expression could be controlled post-transcriptionally. RNA structure predictions using Mfold (Zuker, 2003) identified a putative antiterminator (AT) that overlapped the previously identified terminator (T) hairpin. In addition, a potential anti-antiterminator (AAT) structure was identified even further upstream that overlapped with the AT structure. These structures suggested a model in which formation of the AAT structure would prevent formation of the AT structure, leading to termination upstream of the *yfmG* coding sequence, whereas formation of the AT would lead to transcription readthrough (Fig. 8C, D). Furthermore, inspection of the sequence between the transcription start site and the *yfmG* translation initiation codon led to the identification of a putative 23 amino acid coding sequence (Fig. 8A). This putative minigene was previously annotated as a protein coding sequence (*yfmH*) *yfmH* (Zuber, 2001) and as a sRNA (Irnov et al., 2010). Note that the sRNA is derived from termination at the NusA-dependent terminator (Mondal et al., 2016).

**Figure 8.**
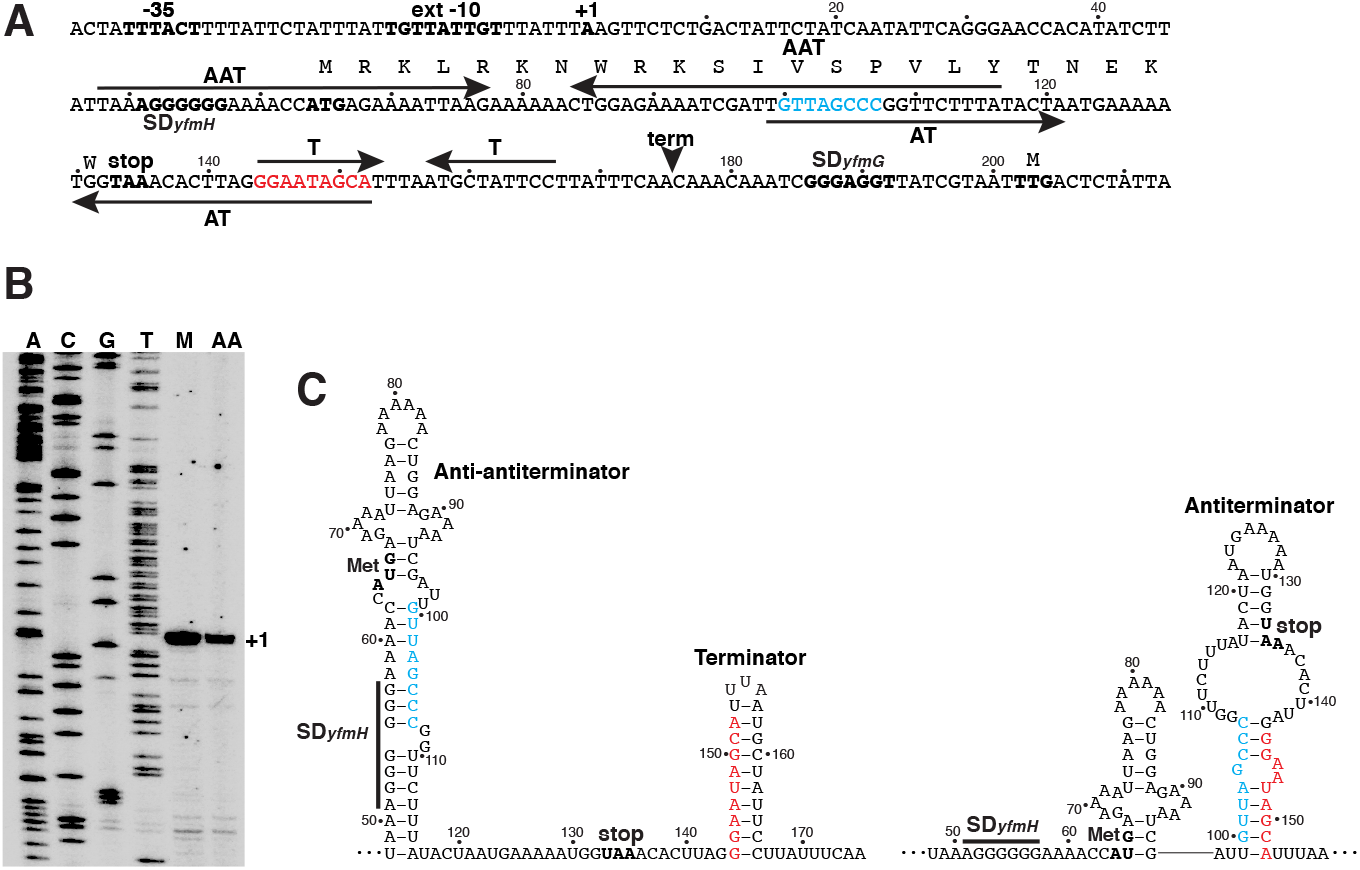
*yfmHG* promoter and leader region. (*A*) Sequence of the *yfmHG* promoter and leader region. The −35 and extended (ext) −10 promoter elements and the transcription start site (+1) are in bold. The *yfmH* Shine-Dalgarno (SD) sequence and the YfmH protein-coding sequence including start (ATG) and stop (TAA) codons are shown. The *yfmG* SD sequence and TTG start codon are also shown. The anti-antiterminator (AAT), antiterminator (AT) and terminator (T) hairpins are indicated by pairs of inverted arrows. The termination site (term) identified in vivo by Term-seq is also marked (Mondal et al., 2016). The overlap between the AAT and AT structures are shown in cyan, while the overlap between the AT and T structures are in red. Numbering is with respect to the start of transcription. (*B*) Primer extension mapping of the *yfmHG* transcription start site. Primer extension analysis was performed on total cellular RNA extracted from a wild type strain of *B. subtilis* grown in minimal + glutamine medium (M) and minimal medium containing all amino acids (AA). The primer extension product identifying the transcription start site is marked (+1). Sequencing lanes (A, C, G, T) are shown. (*C*) Transcription attenuation model. Predicted structures of the anti-antiterminator and terminator structures (left) or the antiterminator structure (right) are shown. Overlaps and color coding between the competing structures are as described in (*A*). The *yfmH* SD sequence, start (met) and stop codons are indicated. Numbering is with respect to the start of transcription.

A chromosomally-integrated *P_yfm_HG-yfmH-lacZ* transcriptional fusion was generated such that the fusion junction was just downstream of the *yfmH* transcription termination site (+175 relative to the start of transcription) (Fusion 1, Fig. 9A). High-level expression was observed for this fusion when grown in the absence of amino acids, whereas expression was about 8-fold lower when grown in the presence of amino acids (Fusion 1, Fig. 9B). These results indicate that *yfmHG* expression is greatly reduced when grown in the presence of amino acids. We next tested a *P_yfm_HG-lacZ* transcriptional fusion in which all but the first 5 nucleotides of the *yfmHG* leader region were deleted (Fusion 2, Fig. 9A). In this case, expression was reduced just 1.5-fold when grown in the presence of amino acids (Fusion 2, Fig. 9B), indicating that amino acids had only a small effect on transcription initiation from the *yfmHG* promoter. A third fusion was generated in which the constitutive *trp* operon promoter was used to replace the *yfmHG* promoter (Fusion 3, Fig. 9A). Expression of this fusion was still strongly reduced (5-fold) when grown in the presence of amino acids (Fusion 3, Fig. 9B). We conclude that amino acid starvation leads to a substantial increase in expression of the *yfmHG* operon and that the long *yfmH* leader but not the promoter is responsible. We further infer that amino acid starvation leads to readthrough past the terminator separating *yfmH* and *yfmG* (Fig. 8).

**Figure 9.**
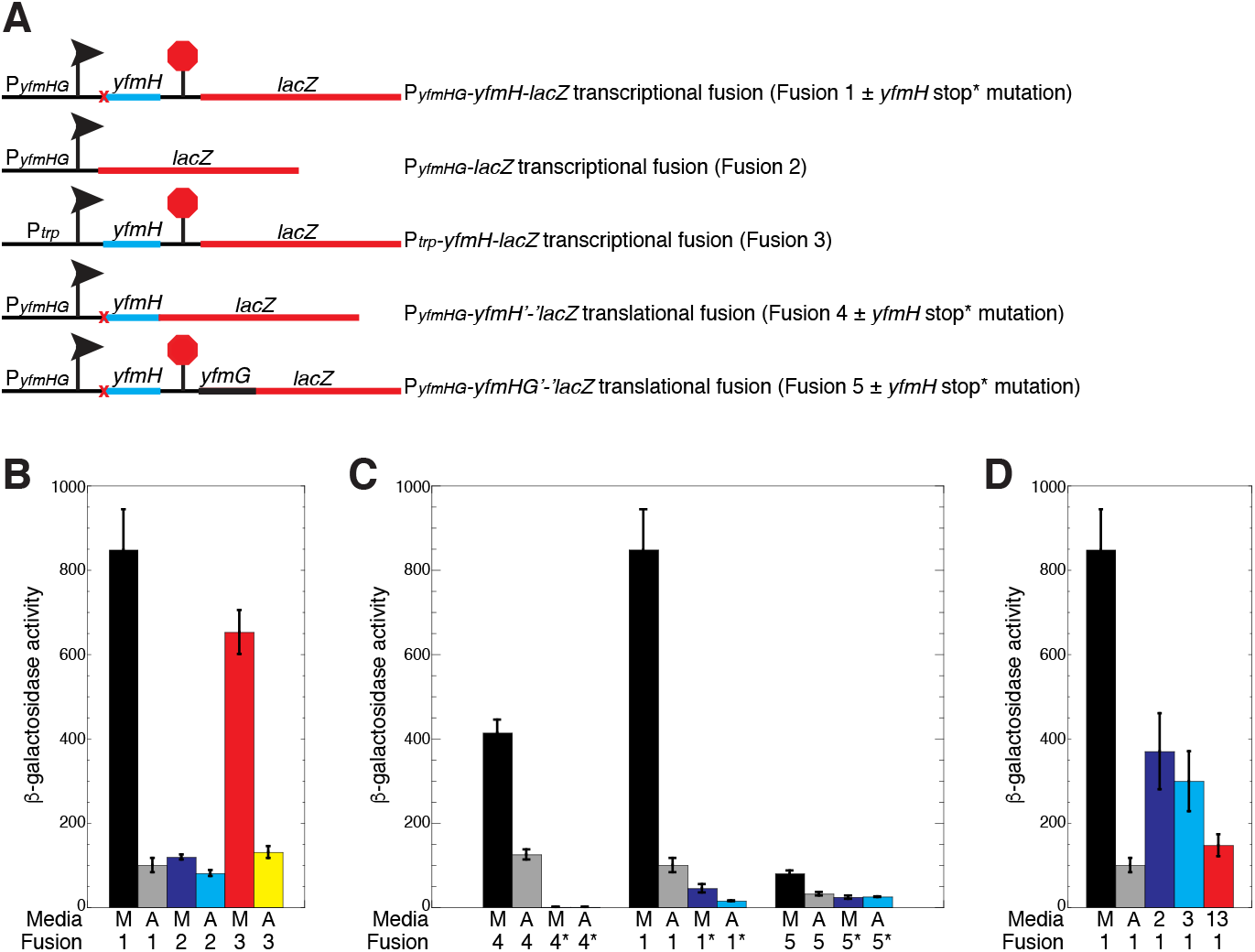
Effects of amino acids and *yfmH* translation on expression of *yfmG*. (*A*) Schematic representation of the *P_yfm_HG-yfmH-lacZ* transcriptional fusion (Fusion 1), *P_yfm_HG-lacZ* transcriptional fusion (Fusion 2), *P_trp_-yfmH-lacZ* transcriptional fusion (Fusion 3), *P_yfm_HG-yfmH’-‘lacZ* translational fusion (Fusion 4), and the *P_yfm_HG-yfmHG’-‘lacZ* translational fusion (Fusion 5). Transcription start site is indicated with an arrow, and the terminator is indicated with a stop sign. *yfmH* start to stop codon mutations are indicated by a red x. (*B-D*) β-galactosidase assays of the indicated chromosomally-integrated fusions when cultures were grown in minimal + glutamine media (M), ACH + tryptophan (A), minimal + lysine and arginine (2), minimal + leucine, isoleucine and valine (3), or minimal + the thirteen amino acids encoded in *yfmH* (13). * indicates a start-to-stop codon mutation in *yfmH*. All cultures contained glutamine. Each experiment was performed at least 3 times. Values are averages ± standard deviation.

A *P_yfm_HG-yfmH’-‘lacZ* translational fusion was generated to determine whether *yfmH* was actually translated (Fusion 4, Fig. 9A). The junction of this fusion was just after the 23^rd^ codon (i.e. the *yfmH* stop codon was replaced with the *lacZ* coding sequence). This fusion was expressed in the absence and presence of amino acids (Fusion 4, Fig. 9C), whereas replacing the AUG start codon of *yfmH* with a UGA stop codon eliminated expression (Fusion 4*, Fig. 9C). These results establish that *yfmH* is translated and likely encodes a leader peptide that regulates *yfmG* expression. We next determined the effect of *yfmH* translation on expression of *P_yfm_HG-yfmH-lacZ* transcriptional and *P_yfm_HG-yfmHG’-‘lacZ* translational fusions (Fusions 1 and 5, Fig. 9A). Introduction of the *yfmH* start to stop codon mutation greatly reduced expression of the transcriptional fusion, however expression of this fusion still responded to amino acids (Fusion 1*, Fig. 9C). Although expression of the *P_yfm_HG-yfmHG’-‘lacZ* translational fusion was low, its expression was reduced 2.5-fold upon addition of amino acids (Fusion 5, Fig. 9C). Importantly, introduction of the *yfmH* start-to-stop codon mutation resulted in 3-fold lower expression and eliminated the effect of amino acids on expression (Fusion 5*, Fig. 9C).

One possibility that could explain this expression data is that the rate of *yfmH* translation would dictate which of the alternative structures form upstream of *yfmG*. Since expression was high in the absence of amino acids, we postulated that a slow rate of *yfmH* translation would favor formation of the AT structure by interfering with formation of the AAT structure (Fig. 8C). Inspection of the *yfmH* coding sequence indicated that it contains codons specifying 13 different amino acids (Fig. 8A). We also noticed that there are three arginine-lysine (RK) motifs in the first half of YfmG, as well as one K further downstream. We found that growth in minimal medium plus just these two amino acids resulted in a 2.5-fold reduction in expression of the *P_yfm_HG-yfmH-lacZ* transcriptional fusion (Fig. 9D). In addition, growth in minimal medium plus isoleucine, leucine and valine (ILV), which corresponds to five YfmH codons, resulted in a 3-fold decrease in expression. Finally, when we grew cultures in minimal medium containing all 13 amino acids encoded in YfmH we found that expression of the *P_yfm_HG-yfmH-lacZ* transcriptional fusion was almost as low as when this strain was grown with all amino acids (Fig. 9D). These results are consistent with a model in which a slow rate of YfmH translation favors formation of the AT structure, leading to transcriptional readthrough into the *yfmG* coding sequence. Conversely, a fast rate of YfmH translation would interfere with AT formation, leading to transcription termination upstream of *yfmG*. This novel type of RNA switch offers a way to simultaneously monitor the levels of several amino acids at once.

## Discussion

RNA structure in bacterial cells plays important and diverse roles on both global and gene-specific levels. Using Structure-seq2 to probe the in vivo RNA structure genome-wide in *B. subtilis* in the presence and absence of amino acids, we saw that there is transcriptome-wide reduced DMS reactivity of RNA in response to amino acid starvation. Similarly, changes in the RNA structurome were observed in eukaryotic species where both heat stress and salinity stress promoted reduced DMS reactivity of RNA (Su et al., 2018; Tack et al., 2020). These findings suggest that there may be a universal stress response mechanism throughout the different domains of life that is manifested in changes in RNA accessibility. One possible explanation for reduced DMS reactivity in the CDS is that translating ribosomes may move more slowly during amino acid starvation, which would passively shield the RNA from DMS. Another explanation is that reduced rates of translation may result in an increase in the distance between translating ribosomes, resulting in the formation of RNA structures that would not have the chance to form when translation is efficient under amino acid replete conditions.

When relating DMS reactivity changes with abundance changes following amino acid starvation, we saw an inverse correlation among all transcripts in which higher average DMS reactivity in the absence of amino acids corresponded to a decrease in transcript abundance. This inverse correlation found in eukaryotes (rice, *Arabidopsis*) (Su et al., 2018; Tack et al., 2020) and now in bacteria (*B. subtilis*) suggests that the regulation of RNA abundance by adjusting transcript accessibility may be a deeply conserved characteristic of all life. Beyond this, the inverse relationship was observed when comparing whole transcripts as well as the CDS, but not when comparing the 5’UTR and 3’UTR regions (Fig. 2B, D). These data suggest that 5’UTRs, which do not contain translating ribosomes nor sequence constraints due to coding, have a wider range of structural change, supporting the notion that most RNA structural changes that regulate gene expression occur within 5’UTRs.

Within *B. subtilis*, the inverse relationship between DMS reactivity and abundance under amino acid starvation conditions is notably pronounced in the stringent response and CodY regulons, both of which are known to be involved in regulating gene expression in response to amino acid starvation. As other regulons did not show this dramatic correlation, these results suggest that only certain transcripts are susceptible to the inverse relationship under these specific conditions, which may provide an advantage to *B. subtilis* under amino acid starvation conditions. We hypothesize that a change in amino acid availability causes direct and indirect effects on the structure of the transcripts within the stringent response and CodY regulons that then causes a corresponding change in abundance. This pattern is likely due to a mixture of mechanisms. For example, during the stringent response, protection from DMS reactivity caused by increased translation would lead to increased transcript abundance by protecting the mRNAs from nucleolytic digestion (Fig. 4A, B teal).

Importantly, we demonstrate that genome-wide RNA structural data are able to report on, as well as predict, specific RNA switches that control gene expression. We showed that under two different conditions of *B. subtilis* growth, Structure-seq2 can accurately characterize RNA switches known to respond to amino acids. While single-gene in vitro characterization can provide detailed mechanistic information on a single riboswitch, in vivo genome-wide studies provide a robust tool to analyze the structure and function of numerous RNA switches in a single study, to do so in the more relevant condition of the cell, and to identify new candidate RNA switches.

By using DMS reactivity values generated through Structure-seq2, we were able to discover novel RNA features through these high throughput sequencing data. Particularly, we identified a transcription attenuation mechanism in the *yfmG* leader region by analyzing regions with high DMS reactivity changes in response to amino acids within the RNA structurome. Within this region, the mRNA encoding a short leader peptide overlaps AAT and AT hairpins, and the absence of amino acids found within this peptide promotes transcription readthrough into *yfmG*. These findings suggest that *yfmG*, a gene with unknown function, may play a role in survival under amino acid starvation conditions. Additional studies are required to elucidate the function of *yfmHG*, and how it relates to starvation for the 13 amino acids encoded in *yfmH*. Uncovering this mechanism has broad implications – it suggests that *B. subtilis* can use a combination of RNA structure and short peptides to sense multiple amino acids. In conclusion, our study demonstrates the utility of using the in vivo genome-wide data generated from Structure-seq2 to discover novel RNA-based regulatory mechanisms.

## Materials And Methods

### Bacterial strains, plasmids and oligonucleotides

*B. subtilis* strains used in this study are listed in Table 1. Plasmids used in this study are described in Supplemental Table M.1 (Grandoni et al., 1993; Merino et al., 1995) and DNA oligonucleotides in Supplemental Table M.2.

**Table 1.**
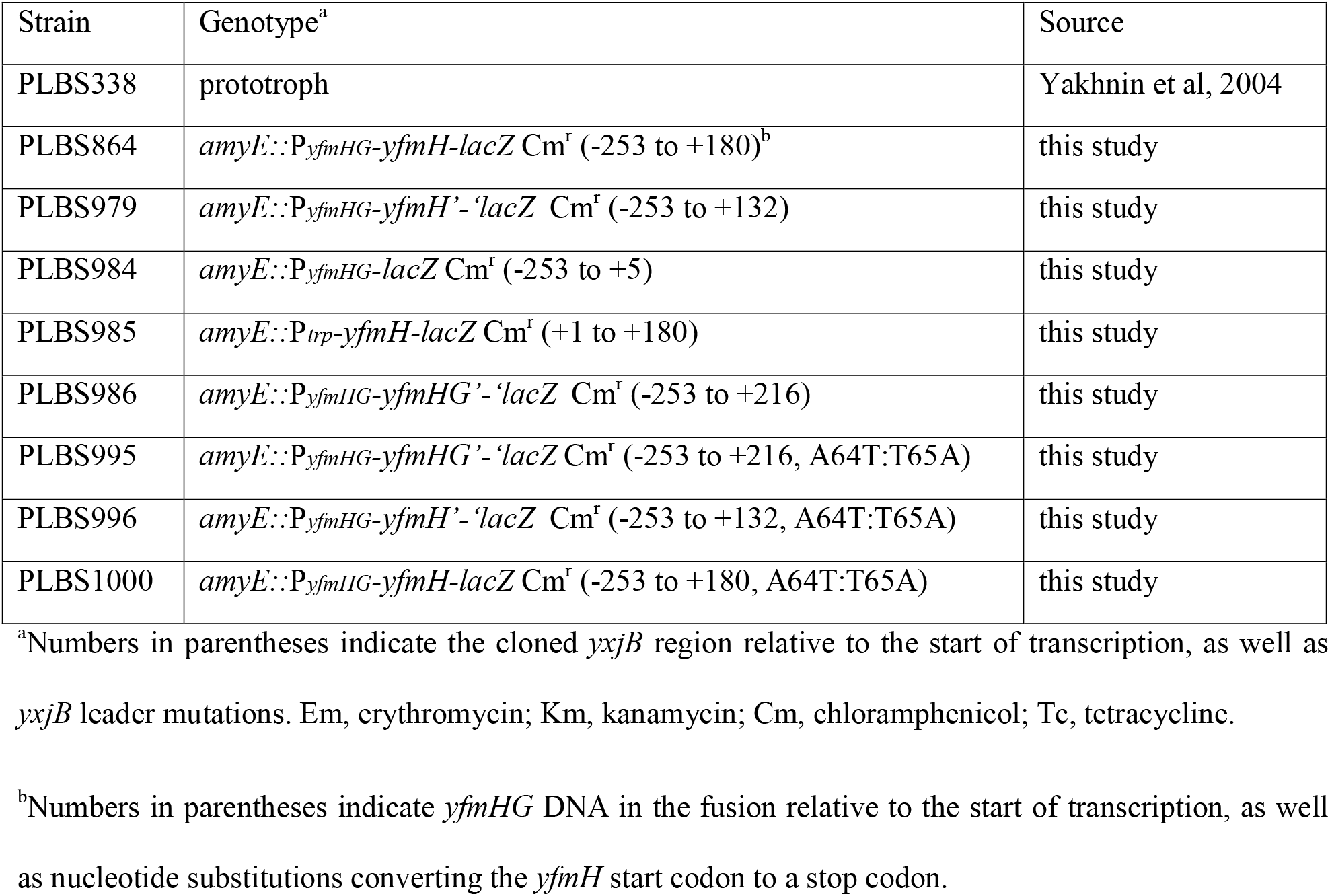
*B. subtilis* strains used in this study.

### Primer extension assay

Total RNA was isolated from late-exponential phase cultures of *B. subtilis* PLBS338 grown in minimal + glutamine (−AA) medium (1X Spizizen salts, 0.5% glucose, 0.04% L-glutamine) and ACH + tryptophan (+AA) medium (1X Spizizen salts, 0.5% glucose, 0.2% acid casein hydrosylate, 0.01% L-tryptophan) using the RNeasy Mini kit (Qiagen). Ten μg of total RNA was hybridized to 150 nM ^32^P 5’-end-labeled DNA oligonucleotide complementary to nucleotides +94 to +109 (relative to *yfmHG* transcription) for 3 min at 80° C. Reaction mixtures (6 μl) containing 2 μl of hybridization mixture, 375 μM of each dNTP, 10 mM dithiothreitol (DTT), 1X SuperScript III buffer, and 0.25μl SuperScript III Reverse Transcriptase (Life Technologies) were incubated for 30 min at 42°C. Reactions were terminated by the addition of 6 μl of stop solution (95% formamide, 20 mM EDTA, 0.025% SDS, 0.025% xylene cyanol, 0.025 % bromophenol blue). Samples were heated for 2 min at 95°C prior to fractionation through standard 6% polyacrylamide sequencing gels. Sequencing reactions were performed using pYH352 as template and the same end-labeled DNA oligonucleotide as a primer. Radiolabeled bands were imaged on a Typhoon 8600 Variable Mode Imager.

### β-galactosidase assay

*B. subtilis* cultures containing various transcriptional or translational fusions were grown at 37°C in −AA medium supplemented with 5 μg/ml chloramphenicol or in +AA medium supplemented with 5 μg/ml chloramphenicol. When indicated, −AA growth media also contained 0.004 % alanine, 0.02 % arginine, 0.004 % asparagine, 0.004 % aspartic acid, 0.002 % cysteine, 0.002 % glutamic acid, 0.01 % glycine, 0.01 % histidine, 0.01 % isoleucine, 0.01 % leucine, 0.008 % lysine, 0.007 % methionine, 0.003 % phenylalanine, 0.02 % proline, 0.01 % serine, 0.01 % threonine, 0.003 % tyrosine, 0.02 % valine and/or 0.01 % L-tryptophan. Cells were grown to mid-exponential phase and β-galactosidase activity was determined as described previously (Du and Babitzke, 1998).

### Bacterial growth and DMS treatment

DMS treatments were performed in a chemical fume hood. *B. subtilis* strain PLBS338 (Yakhnin et al., 2004) was grown at 37 °C in +AA or −AA media. For each replicate (three per growth media), cultures were grown to mid-exponential phase prior to DMS treatment. For the -DMS control, 6 mL of culture was transferred to a tube containing 0.45 g DTT and slightly agitated for 2 min. The solution was then poured into an equal volume of frozen slurry killing buffer (10 mM Tris-HCl, pH 7.2, 5 mM MgCl_2_, 25 mM NaN_3_, 1.5 mM chloramphenicol, and 12.5% ethanol. DMS was added to the remaining 14 mL of culture to a final concentration of 50 mM and the reaction was allowed to proceed for 5 min in a 37 °C in a shaking water bath. These conditions provide appropriate single-hit kinetics (Supplemental Fig. S1). After 5 min, 1 g DTT was added to the culture to quench the reaction for 2 min in the shaking water bath. 6 mL of the quenched culture was then added to frozen slurry killing buffer. Culture tubes were centrifuged for 10 min at 10,000 rpm. Cell pellets were suspended in 1 mL ice cold killing buffer, and then cells were pelleted in a microfuge tube. The wash procedure was repeated and then cell pellets were stored at −20 °C.

### Structure-seq2 library preparation

To obtain RNA for Structure-seq2 library preparation, total RNA from the cell pellets was extracted using the RNeasy Mini Kit (Qiagen). After extraction, the quality of RNA was assessed by observing rRNA bands following fractionation through a 1% agarose gel, as well as by a Prokaryote Total RNA Nano Bioanalyzer scan to identify rRNA peaks. rRNA was then depleted using the Ribo-Zero rRNA Removal Kit for Gram-positive bacteria (Illumina), and the yield and quality were assessed via spectrophotometry and by an mRNA Pico Bioanalyzer scan to confirm depletion of the rRNA peaks.

Following rRNA depletion, RNA was either sent for RNA-seq analysis or libraries were prepared following the biotin variation of Structure-seq2, as described previously (Ritchey et al., 2019). Briefly for Structure-seq2 preparation, using ~300 ng rRNA-depleted RNA, cDNA was synthesized using a random hexamer to prime reverse transcription in the presence of 500 μM each dNTP, 62.5 μM biotinylated-dCTP, and 62.5 μM biotinylated-dUTP. RT stops immediately before the site of DMS modification, allowing these modifications to be read out transcriptome-wide. The reactions were purified via RNA Clean & Concentrator-5 (Zymo Research) and streptavidin pull-down. A hairpin adaptor was then ligated to the 3’ end of the cDNA fragments, and products were purified via RNA Clean & Concentrator-5 and streptavidin pull down. A PCR cycle test was performed to determine the appropriate number of cycles for amplification and 18 cycles was chosen. PCR was performed with sequences complementary to Illumina sequencing primers incorporated into the primers. The resulting libraries were fractionated through a denaturing 8.3 M urea 10% polyacrylamide gel, purified via a crush and soak method to select for sizes 200-600 nt, and submitted for sequencing. Prior to sequencing, the quality of libraries was assessed via Bioanalyzer scans.

### Transcriptome Assembly

To maximize the amount of resolvable sequence space containing potential structural regulatory elements (i.e. 5’ and 3’ UTRs), we generated our own reference transcriptome. Standard RNA-seq libraries were generated from both +AA and −AA conditions, yielding 39,310,106 and 42,864,834 reads respectively. These libraries were trimmed with trimmomatic, version 0.36 (Bolger et al., 2014) yielding 39,280,081 +AA and 41,830,095 −AA reads for assembly. Trimmed reads were used as input for Rockhopper, version 2.03 (Tjaden, 2015), which aligned 38,256,615 and 40,582,875 reads on the *B. subtilis* str. 168 genome for a combined assembly containing 2,369 5’UTR regions and 2,119 3’UTR regions. The identified boundaries of these transcripts were used to parse out the putative transcripts (https://github.com/StructureFold2/NotToosubtilis) into fasta format. The fasta file of this whole transcriptome is included in Supplemental Item S2.

To confirm the efficacy of our Structure-Seq2/StructureFold2 method in finding RNA switches, we manually generated a panel of the complete transcripts of known riboswitches, thus relying on annotation where it was available, rather than relying on a reference-based assembly as with the whole transcriptome. Sequences were found with BsubCyc (Caspi et al., 2014), and compiled manually into a fasta file. Where transcription start sites were not noted in BsubCyc, they were estimated based on the beginning of the annotated regulatory features (e.g. T-box) upstream of the gene. The fasta file of this targeted transcriptome is included in Supplemental Item S2.

### Reactivity analyses

Structure-Seq2 libraries were first trimmed via Cutadapt, version 1.16 (Martin, 2011) via the fastq_trimmer module of StructureFold2 (Tack et al., 2018) (Supplemental Table W.1). These trimmed reads were aligned to both the whole transcriptome (for all genome-wide analyses) and to the panel of known riboswitches (for targeted analyses) via Bowtie2, version 2.3.4.1 (Langmead and Salzberg, 2012) via the fastq_mapper module of StructureFold2 (Tack et al., 2018) (Supplemental Table W.2.a, W.2.b). Both sets of mappings were filtered to remove reads with a mismatch on the first base or reads with more than 3 mismatches with the StructureFold2 sam_filter module (Tack et al., 2018) (Supplemental Table W.3.a and W.3.b) which utilizes SAMtools, version 0.1.19 (Li et al., 2009).

Filtered mappings were converted to .rtsc (reverse transcriptase stop count) files, replicates were merged, and per transcript coverage was calculated via StructureFold2 (Tack et al., 2018). For transcripts above the coverage threshold of 1 in the merged file (amounting to ≥1 RT stop per A or C residue in the transcript) (Ding et al., 2014; Su et al., 2018; Tack et al., 2020), we correlated raw RT stop patterns between component replicates (Supplemental Fig. S3; Supplemental Table T.1) with R (R Core Team, 2019) using the plyr package (Wickham, 2011) via the rtsc_correlation module of StructureFold2 (Tack et al., 2018); this follows the same process for evaluating library replicate consistency as was done previously (Su et al., 2018; Tack et al., 2020; Waldron et al., 2019). Merged rtsc files were then used to calculate per-base reactivity via the rtsc_to_react module of StructureFold2 (Tack et al., 2018), yielding react files.

React files for the whole transcriptome analyses were analyzed with the StructureFold2 (Tack et al., 2018) react_statistics module to obtain overall transcript reactivity statistics. To estimate transcript abundance, the corresponding +AA/-AA rtsc files were used to calculate transcripts per million reads (TPM) (Wagner et al., 2012) via the StructureFold2 (Tack et al., 2018) rtsc_abundance module. The DMS reactivity statistics and abundances were analyzed in R (Team, 2008), and plotted using the ggplot package (Wickham, 2016). Fisher Z-tests between correlations were calculated with the cocor package (Diedenhofen and Musch, 2015). For batch analysis comparing our data to know regulons, we integrated a list of regulons from *SubtiWiki* (Zhu and Stulke, 2018) (http://www.subtiwiki.uni-goettingen.de/v3/exports). For the sliding window analyses, we used the react_windows module (Tack et al., 2018) to find areas of reactivity change between conditions. We queried by net change (delta) and searched for the top 5% of most changed 20 nt windows sliding by 10 nt within the 5’UTRs of all transcripts (Supplemental Table L.6).

The RT stop counts of the targeted switch transcript 5’UTRs were first subset from the entire targeted switch transcript counts, creating separate 5’UTR exclusive rtsc files. These rtsc files were then used to calculate per base reactivity on the 5’UTR in both +AA and −AA conditions, applying a 5’UTR specific 2-8% scale from +AA to the −AA data, thus generating react files. These react files containing the final calculated reactivity values were then converted to .csv files (included in Supplemental Item S3) and manually analyzed for changes, or used as restraints to guide folding using RNAStructure (Reuter and Mathews, 2010) via StructureFold2 (Tack et al., 2018). For the sliding window analysis on the targeted windows, we followed a similar procedure as above and queried for gains, losses, and net change. The fasta version of these results were submitted for MEME analysis using the online webserver with default parameters (Bailey et al., 2009). Only the top 25% of windows that gained reactivity produced significant motif enrichment (Supplemental Table L.5; Fig. 6E, F). Structure-Seq2 and RNA-seq Libraries were deposited in Gene Expression Omnibus (GEO) (accession number GSE148936). The reviewer access code is mhadkuksvdmhhmz.

### Library quality analysis

We used StructureFold2 (Tack et al., 2018) to process and analyze the resulting Structure-seq2 libraries. The rtsc_correlation.py module was used to identify the RT stops prior to every A and C residue in our assembled transcriptome in .csv format for all +DMS libraries in both treatments (+AA, −AA). These two .csv files were read in and analyzed using R (Team, 2008) and the ggplot2 package (Wickham, 2016). For every pairwise combination of two replicates from the same treatment, we treated the entire transcriptome as one entity and computed the correlation coefficient of the number of RT stops between every A and C: values are included on each respective graph and in Supplemental Table T.1. Second, we subdivided the transcriptome by transcript and computed the correlation coefficient for each transcript between each pairwise combination of two replicates. The density of these distributions of r values were likewise plotted, showing the vast majority of per transcript correlation coefficients to be high (Supplemental Fig. S3; Supplemental Table T.1). These results implied that the replicates of both treatments were suitably similar to be pooled to minimize technical variation and provide more robust numbers before proceeding to calculate derived DMS reactivity.

## Supplemental Information

1. Supplemental Figures S1-S5 (figures, legends, and references
2. Supplemental Item S1 (extended regulon figure)
3. Supplemental Item S2 (transcriptome assembly information)
4. Supplemental Item S3 (reactivity value information)
5. Supplemental Tables (M.1, M.2, W.1-W.4, T.1-T.4, D.1-D.10, L.1-L.6)

## Acknowledgments

This work was supported by National Institutes of Health grants R01-GM098399 to P.B., R35-GM127064 to P.C.B, and F32-GM128311 to E.A.J., National Science Foundation grant IOS-1339282 to P.C.B. and S.M.A., and by grant KA2016-85222 from the Charles E. Kaufman Foundation of the Pittsburgh Foundation to P.B., P.C.B. and S.M.A.

## REFERENCES

Ames TD, Breaker RR. 2011. Bacterial aptamers that selectively bind glutamine. RNA Biol 8: 82–89.

Babitzke P, Baker C., Romeo T. 2009. Regulation of translation initiation by RNA binding proteins. Annu Rev Microbiol 63: 27–44.

Babitzke P, Gollnick P. 2001. Posttranscription initiation control of tryptophan metabolism in *Bacillus subtilis* by the *trp* RNA-binding attenuation protein (TRAP), anti-TRAP, and RNA structure. J Bacteriol 183:5795–5802.

Bailey TL, Boden M, Buske FA, Frith M, Grant CE, Clementi L, Ren J, Li WW, Noble WS. 2009. MEME SUITE: tools for motif discovery and searching. Nucleic Acids Res 37: W202–208.

Bolger AM, Lohse M, Usadel B. 2014. Trimmomatic: a flexible trimmer for Illumina sequence data. Bioinformatics 30: 2114–2120.

Brinsmade SR, Alexander EL, Livny J, Stettner AI, Segre D, Rhee KY, Sonenshein AL. 2014. Hierarchical expression of genes controlled by the *Bacillus subtilis* global regulatory protein CodY. Proc Natl Acad Sci U S A 111: 8227–8232.

Burkhardt DH, Rouskin S, Zhang Y, Li GW, Weissman JS, Gross CA. 2017. Operon mRNAs are organized into ORF-centric structures that predict translation efficiency. Elife 6.

Carlson BL, Ballister ER, Skordalakes E, King DS, Breidenbach MA, Gilmore SA, Berger JM, Bertozzi CR. 2008. Function and structure of a prokaryotic formylglycine-generating enzyme. J Biol Chem 283: 20117–20125.

Caserta E, Liu LC, Grundy FJ, Henkin TM. 2015. Codon-anticodon recognition in the *Bacillus subtilis glyQS* T box riboswitch: RNA-dependent codon selection outside the ribosome. J Biol Chem 290: 23336–23347.

Caspi R, Altman T, Billington R, Dreher K, Foerster H, Fulcher CA, Holland TA, Keseler IM, Kothari A, Kubo A., et al. 2014. The MetaCyc database of metabolic pathways and enzymes and the BioCyc collection of Pathway/Genome Databases. Nucleic Acids Res 42: D459–471.

Cavanagh AT Wassarman KM. 2014. 6S RNA, a global regulator of transcription in *Escherichia coli, Bacillus subtilis*, and beyond. Annu Rev Microbiol 68: 45–60.

Del Campo C, Bartholomaus A, Fedyunin I, Ignatova Z. 2015. Secondary structure across the bacterial transcriptome reveals versatile roles in mRNA regulation and function. PLoS Genet 11: e1005613.

Diedenhofen B, Musch J. 2015. cocor: a comprehensive solution for the statistical comparison of correlations. PLoS One 10: e0121945.

Ding Y, Tang Y, Kwok CK, Zhang Y, Bevilacqua PC, Assmann SM. 2014. In vivo genome-wide profiling of RNA secondary structure reveals novel regulatory features. Nature 505: 696–700.

Du H, Babitzke P. 1998. *trp* RNA-binding attenuation protein-mediated long distance RNA refolding regulates translation of *trpE* in Bacillus subtilis. J Biol Chem 273: 20494–20503.

Eymann C, Homuth G, Scharf C, Hecker M. 2002. *Bacillus subtilis* functional genomics: global characterization of the stringent response by proteome and transcriptome analysis. J Bacteriol 184: 2500–2520.

Eymann C, Mittenhuber G, Hecker M. 2001. The stringent response, sigmaH-dependent gene expression and sporulation in *Bacillus subtilis*. Mol Gen Genet 264: 913–923.

Fujita Y. 2009. Carbon catabolite control of the metabolic network in *Bacillus subtilis*. Biosci Biotechnol Biochem 73: 245–259.

Geiger T, Wolz C. 2014. Intersection of the stringent response and the CodY regulon in low GC Grampositive bacteria. Int J Med Microbiol 304: 150–155.

Gollnick P,l Babitzke P. 2002. Transcription attenuation. Biochim Biophys Acta 1577: 240–250.

Gollnick P, Babitzke P, Antson A, Yanofsky C. 2005. Complexity in regulation of tryptophan biosynthesis in *Bacillus subtilis*. Annu Rev Genet 39: 47–68.

Grandoni JA, Fulmer SB, Brizzio V, Zahler SA, Calvo JM. 1993. Regions of the *Bacillus subtilis ilv-leu* operon involved in regulation by leucine. J Bacteriol 175: 7581–7593.

Green NJ, Grundy FJ, Henkin TM. 2010. The T box mechanism: tRNA as a regulatory molecule. FEBS Lett 584: 318–324.

Grigg JC, Ke A. 2013. Structural determinants for geometry and information decoding of tRNA by T box leader RNA. Structure 21: 2025–2032.

Grundy FJ, Henkin TM. 1998. The S box regulon: a new global transcription termination control system for methionine and cysteine biosynthesis genes in gram-positive bacteria. Mol Microbiol 30: 737–749.

Grundy FJ, Henkin TM. 2004. Regulation of gene expression by effectors that bind to RNA. Curr Opin Microbiol 7: 126–131.

Grundy FJ, Lehman SC, Henkin TM. 2003. The L box regulon: lysine sensing by leader RNAs of bacterial lysine biosynthesis genes. Proc Natl Acad Sci U S A 100: 12057–12062.

Hecker M, Volker U. 2001. General stress response of *Bacillus subtilis* and other bacteria. Adv Microb Physiol 44: 35–91.

Irnov I, Sharma CM, Vogel J, Winkler WC. 2010. Identification of regulatory RNAs in *Bacillus subtilis*. Nucleic Acids Res 38: 6637–6651.

Kovacs AT. 2016. Bacterial differentiation via gradual activation of global regulators. Curr Genet 62: 125–128.

Langmead B, Salzberg SL. 2012. Fast gapped-read alignment with Bowtie 2. Nat Methods 9: 357–359.

Li F, Zheng Q, Vandivier LE, Willmann MR, Chen Y, Gregory BD. 2012. Regulatory impact of RNA secondary structure across the *Arabidopsis* transcriptome. Plant Cell 24: 4346–4359.

Li H, Handsaker B, Wysoker A, Fennell T, Ruan J, Homer N, Marth G, Abecasis G, Durbin R; 1000 Genome Project Data Subgroup. 2009. The sequence alignment/map format and SAMtools. Bioinformatics 25: 2078–2079.

Lu C, Ding F, Chowdhury A, Pradhan V, Tomsic J, Holmes WM, Henkin TM, Ke A. 2010. SAM recognition and conformational switching mechanism in the *Bacillus subtilis yitJ* S box/SAM-I riboswitch. J Mol Biol 404: 803–818.

Mandal M, Lee M, Barrick JE, Weinberg Z, Emilsson GM, Ruzzo WL, Breaker RR. 2004. A glycine-dependent riboswitch that uses cooperative binding to control gene expression. Science 306: 275–279.

Martin M. 2011. Cutadapt removes adapter sequences from high-throughput sequencing reads. EMBnet Journal 17:10–12.

McCown PJ, Corbino KA, Stav S, Sherlock ME, Breaker RR. 2017. Riboswitch diversity and distribution. RNA 23: 995–1011.

McDaniel BA, Grundy FJ, Artsimovitch I, Henkin TM. 2003. Transcription termination control of the S box system: direct measurement of S-adenosylmethionine by the leader RNA. Proc Natl Acad Sci U S A 100:3083–3088.

Merino E, Babitzke P, Yanofsky C. 1995. *trp* RNA-binding attenuation protein (TRAP)-*trp* leader RNA interactions mediate translational as well as transcriptional regulation of the *Bacillus subtilis trp* operon. J Bacteriol 177: 6362–6370.

Mondal S, Yakhnin AV, Sebastian A, Albert I, Babitzke P. 2016. NusA-dependent transcription termination prevents misregulation of global gene expression. Nat Microbiol 1: 15007.

Mustoe AM, Busan S, Rice GM, Hajdin CE, Peterson BK, Ruda VM, Kubica N, Nutiu R, Baryza JL, Weeks KM. 2018. Pervasive Regulatory Functions of mRNA Structure Revealed by High-Resolution SHAPE Probing. Cell 173: 181-195.e118.

R Core Team. 2019. R: A language and enviornment for statistical computing. R Foundation for Statistical Computing, Vienna, Austria. URL https://www.R-project.org/.

Reuter JS, Mathews DH. 2010. RNAstructure: software for RNA secondary structure prediction and analysis. BMC Bioinformatics 11: 129.

Righetti F, Nuss AM, Twittenhoff C, Beele S, Urban K, Will S, Bernhart SH, Stadler PF, Dersch P, Narberhaus F. 2016. Temperature-responsive in vitro RNA structurome of *Yersinia pseudotuberculosis*. Proc Natl Acad Sci US A 113: 7237–7242.

Ritchey LE, Su Z, Assmann SM, Bevilacqua PC. 2019. In vivo genome-wide RNA structure probing with Structure-seq. Methods Mol Biol 1933: 305–341.

Ritchey LE, Su Z, Tang Y, Tack DC, Assmann SM, Bevilacqua PC. 2017. Structure-seq2: sensitive and accurate genome-wide profiling of RNA structure in vivo. Nucleic Acids Res 45: e135.

Romeo T, Vakulskas CA, Babitzke P. 2013. Post-transcriptional regulation on a global scale: form and function of Csr/Rsm systems. Environ Microbiol 15: 313–324.

Rouskin S, Zubradt M, Washietl S, Kellis M, Weissman JS. 2014. Genome-wide probing of RNA structure reveals active unfolding of mRNA structures in vivo. Nature 505: 701–705.

Sherwood AV, Henkin TM. 2016. Riboswitch-mediated gene regulation: Novel RNA architectures dictate gene expression responses. Annu Rev Microbiol 70: 361–374.

Sonenshein AL. 2005. CodY, a global regulator of stationary phase and virulence in Gram-positive bacteria. Curr Opin Microbiol 8: 203–207.

Stav S, Atilho RM, Mirihana Arachchilage G, Nguyen G, Higgs G, Breaker RR. 2019. Genome-wide discovery of structured noncoding RNAs in bacteria. BMC Microbiol 19: 66.

Su Z, Tang Y, Ritchey LE, Tack DC, Zhu M, Bevilacqua PC, Assmann SM. 2018. Genome-wide RNA structurome reprogramming by acute heat shock globally regulates mRNA abundance. Proc Natl Acad Sci USA 115: 12170–12175.

Sudarsan N, Wickiser JK, Nakamura S, Ebert MS, Breaker RR. 2003. An mRNA structure in bacteria that controls gene expression by binding lysine. Genes Dev 17: 2688–2697.

Tack DC, Su Z, Yu Y, Bevilacqua PC, Assmann SM. 2020. Tissue-specific changes in the RNA structurome mediate salinity response in Arabidopsis. RNA 26: 492–511.

Tack DC, Tang Y, Ritchey LE, Assmann SM, Bevilacqua PC. 2018. StructureFold2: Bringing chemical probing data into the computational fold of RNA structural analysis. Methods 143: 12–15.

Tapsin S, Sun M, Shen Y, Zhang H, Lim XN, Susanto TT, Yang SL, Zeng GS, Lee J, Lezhava A, et al. 2018. Genome-wide identification of natural RNA aptamers in prokaryotes and eukaryotes. Nat Commun 9: 1289.

Tjaden B. 2015. De novo assembly of bacterial transcriptomes from RNA-seq data. Genome Biol 16: 1.

Voelker U, Voelker A, Maul B, Hecker M, Dufour A, Haldenwang WG. 1995. Separate mechanisms activate sigma B of *Bacillus subtilis* in response to environmental and metabolic stresses. J Bacteriol 177: 3771–3780.

Wagner GP, Kin K, Lynch VJ. 2012. Measurement of mRNA abundance using RNA-seq data: RPKM measure is inconsistent among samples. Theory Biosci 131: 281–285.

Waldron JA, Tack DC, Ritchey LE, Gillen SL, Wilczynska A, Turro E, Bevilacqua PC, Assmann SM, Bushell M, Le Quesne J. 2019. mRNA structural elements immediately upstream of the start codon dictate dependence upon eIF4A helicase activity. Genome Biol 20: 300.

Waters LS, Storz G. 2009. Regulatory RNAs in bacteria. Cell 136: 615–628.

Weinberg Z, Lunse CE, Corbino KA, Ames TD, Nelson JW, Roth A, Perkins KR, Sherlock ME, Breaker RR. 2017. Detection of 224 candidate structured RNAs by comparative analysis of specific subsets of intergenic regions. Nucleic Acids Res 45: 10811–10823.

Wickham H. 2011. The split-apply-combine strategy for data analysis. J Stat Softw 40: 29.

Wickham H. 2016. ggplot2: Elegant graphics for data analysis. Springer-Verlang New York, NY.

Winkler WC, Nahvi A, Sudarsan N, Barrick JE, Breaker RR. 2003. An mRNA structure that controls gene expression by binding S-adenosylmethionine. Nat Struct Biol 10: 701–707.

Yakhnin H, Zhang H, Yakhnin AV, Babitzke P. 2004. The *trp* RNA-binding attenuation protein of *Bacillus subtilis* regulates translation of the tryptophan transport gene *trpP (yhaG)* by blocking ribosome binding. J Bacteriol 186: 278–286.

Zhang J, Ferre-D’Amare AR. 2013. Co-crystal structure of a T-box riboswitch stem I domain in complex with its cognate tRNA. Nature 500: 363–366.

Zhu B, Stulke J. 2018. SubtiWiki in 2018: from genes and proteins to functional network annotation of the model organism *Bacillus subtilis*. Nucleic Acids Res 46: D743–D748.

Zuber P. 2001. A peptide profile of the *Bacillus subtilis* genome. Peptides 22: 1555–1577.

Zuker M. 2003. Mfold web server for nucleic acid folding and hybridization prediction. Nucleic Acids Res 31:3406–3415.

